# Generation of a DAT-Flp mouse line for intersectional genetic targeting of dopamine neuron subpopulations

**DOI:** 10.1101/2020.06.24.167908

**Authors:** Daniel J. Kramer, Polina Kosillo, Drew Friedmann, David Stafford, Liqun Luo, Angus Yiu-Fai Lee, Dirk Hockemeyer, John Ngai, Helen S. Bateup

**Author notes:** National Institute of Neurological Disorders and Stroke, National Institutes of Health, Bethesda, MD, USA. Corresponding author: Helen S. Bateup, University of California, Berkeley, 142 Weill Hall (LSA), Berkeley, CA 94720-3200.

## Abstract

Dopamine neurons project to diverse regions throughout the brain to modulate various brain processes and behaviors. It is increasingly appreciated that dopamine neurons are heterogeneous in their gene expression, circuitry, physiology, and function. Current approaches to target dopamine neurons are largely based on single gene drivers, which either label all dopamine neurons, or mark a sub-set but concurrently label non-dopaminergic neurons. Here we establish a novel mouse line in which Flp recombinase is knocked-in to the endogenous *Slc6a3* (dopamine active transporter, DAT) locus. DAT-Flp mice can be used with various Cre-expressing mouse lines to efficiently and selectively label dopaminergic subpopulations using Cre/Flp-dependent intersectional strategies. We demonstrate the utility of this approach by crossing DAT-Flp mice with NEX-Cre mice, to specifically label *Neurod6*-expressing dopamine neurons that project to the nucleus accumbens medial shell. DAT-Flp mice represent a novel tool, which will help parse the diverse functions mediated by dopaminergic circuits.

## Introduction

Despite their modest numbers, dopamine (DA) neurons project widely throughout the brain and influence a diverse set of behaviors and neural processes including movement, motivation, reward learning, and cognition (1,2). Historically, DA neurons have been categorized into 17 main sub-types, the cell groups A1-A17 (1,3). A9 and A10 are the most numerous and reside in the midbrain in the substantia nigra pars compacta (SNc) and ventral tegmental area (VTA), respectively. It has long been recognized that SNc and VTA DA neurons differ significantly in their projection targets, electrophysiological properties, behavioral influence, and vulnerability to degeneration (1,4-11). Recent single cell profiling studies have further revealed that the SNc and VTA are, in fact, made up of smaller, partially overlapping populations based on their gene expression patterns (12-17). How these dopaminergic subpopulations differ in terms of their anatomical location, inputs and outputs, neurotransmitter identity and functional properties is just beginning to be defined.

To better understand the connections and behavioral functions of newly identified DA neuron subpopulations, additional tools are needed that allow genetic access to these subtypes. This is essential as even neighboring DA neurons can project to different places in the brain and exert opposing behavioral effects, i.e. signaling reward versus aversion (8,9). From gene expression profiling studies, it is clear that combinatorial approaches utilizing the intersectional expression of two or more genes will be needed as there are rarely single marker genes that are exclusively expressed by a given subtype (17). In addition, even when the expression of a single gene can distinguish discrete DA subpopulations, these genes may also be expressed in non-dopaminergic neurons and are thus not useful for exclusive targeting of DA neuron subtypes (12).

To tackle this challenge, intersectional mouse genetic approaches have been developed whereby the combinatorial expression of different recombinases can be used to define a neuronal subpopulation (18-21). This approach was recently applied to the DA system through the generation of mice that express Flp recombinase from the endogenous tyrosine hydroxylase (*Th*) locus (18). Using these Th-2A-Flpo mice, Poulin and colleagues successfully generated detailed maps of the projection targets of various genetically-defined dopaminergic subpopulations. However, TH is the rate limiting enzyme in the synthesis of all catecholamines, including DA, norepinephrine and epinephrine. Thus, available transgenic mouse lines that use *Th* to drive expression of recombinases and reporter genes in DA neurons (e.g. Th-IRES-Cre, Th-2A-Flpo, and TH-EGFP) also have expression in other catecholamine-expressing neurons (18,22-25). As a result, the use of *Th*-driven reporters to investigate the behavioral functions of specific dopaminergic circuits may be confounded by contributions from these other cell types (22).

To address this limitation and provide an additional tool for studying dopaminergic subpopulations, we generated a mouse that expresses Flp recombinase from the endogenous *Slc6a3* (dopamine active transporter, DAT) locus. DAT is responsible for the re-uptake of DA back into the presynaptic terminal following release and is expressed almost exclusively in dopaminergic neurons (3,22,23). As a result, DAT-reporter mouse lines have been shown to be more selective and label canonical DA neurons more specifically than *Th*-driven lines (22,26,27). Here we show that midbrain DA neurons are robustly and selectively labeled in DAT-Flp mice using Flp-dependent viral or transgenic reporters.

To provide proof-of-concept for intersectional labeling of a DA neuron subpopulation, we crossed DAT-Flp mice with NEX-Cre mice (28) and verified selective targeting of *Neurod6*-positive DA neurons in the ventromedial VTA. We further performed Adipo-Clear (29) brain clearing and fast scan cyclic voltammetry to demonstrate the highly selective anatomical and functional output of this discrete DA neuron subpopulation. DAT-Flp mice therefore represent a novel tool, which adds to the growing arsenal of mouse genetic approaches that allow precise targeting of specific DA neuron subpopulations. Collectively, these new mouse models provide a valuable means to dissect the contributions of dopaminergic sub-circuits to behavior.

## Results

### Generation of DAT-Flp mice

To allow for intersectional targeting of dopaminergic subpopulations, we developed a DAT-Flp mouse line that expresses Flp recombinase from the endogenous *Slc6a3* (DAT) locus. We used CRISPR/Cas9-based gene editing to knock-in a P2A-Flpo-recombinase sequence 3’ to the final coding exon of DAT, replacing the stop codon in the 15^th^ exon and the 3’ UTR (Fig. 1A). The self-cleaving P2A sequence allows for bicistronic expression of both endogenous DAT protein and Flp recombinase following translation of a single mRNA. For gene editing, we used mouse embryonic stem cells derived from the Ai65(RCFL-tdT)-D mouse line (herein referred to as Ai65) (30). These cells harbor a Flp- and Cre-dependent tdTomato fluorescent reporter in the Rosa26 locus. F0 chimeric mice were PCR genotyped to confirm successful targeting (Fig. 1B). F0 male mice had successful germline transmission and produced pups that expressed the Flp recombinase insert (Fig. 1B).

**Figure 1.**
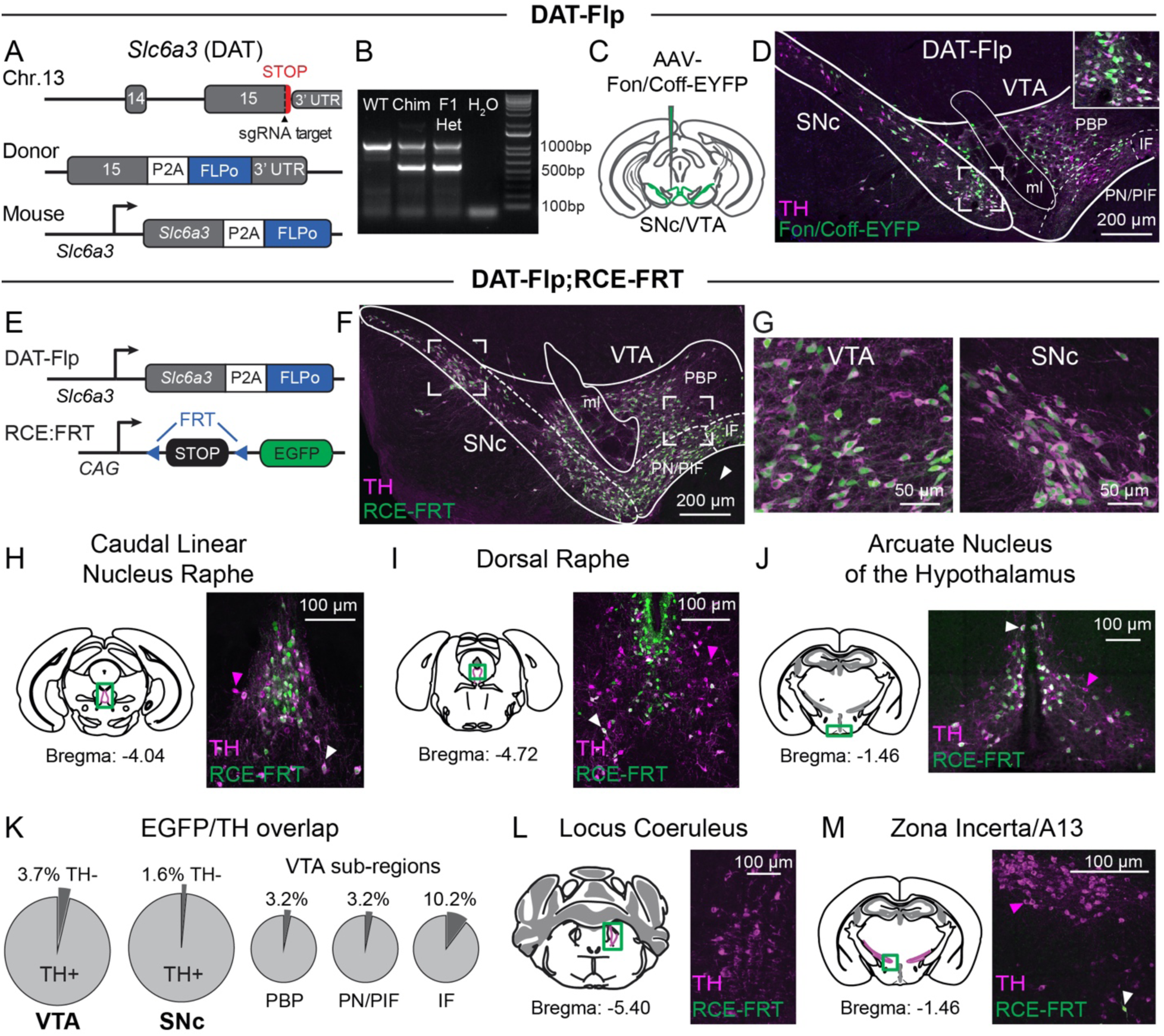
Generation of DAT-Flp mice. A) Targeting scheme to generate DAT-Flp knock-in mice. Top: shown are the last two exons of the *Slc6a3* (DAT) gene on chromosome 13 and the position of the sgRNA. Middle: a targeting donor was used to replace the STOP codon of *Slc6a3* with a P2A sequence followed by the Flpo recombinase reading frame. Bottom: correct targeting leads to a modified locus expressing Flpo under the control of the endogenous *Slc6a3* regulatory elements. B) PCR genotyping strategy for DAT-Flp mice. The 1000 bp band denotes the WT allele. The 650 bp band indicates the presence of the P2A-Flpo insert. Expression in the F1 generation indicates successful germline transmission. Chim = chimeric founders (F0). C) Schematic of the unilateral injection of AAV-Fon/Coff-EYFP into the medial substantia nigra pars compacta (SNc) and ventral tegmental area (VTA). D) Coronal midbrain section stained with an antibody against tyrosine hydroxylase (TH, magenta) showing expression of Fon/Coff-EYFP in the midbrain (green). Inset shows a zoomed-in image of SNc neurons. PBP = parabrachial pigmented nuclei; PN/PIF = paranigral nucleus/parainterfascicular nucleus; IF = interfascicular nucleus; ml = medial lemniscus. E) Schematic of the genetic cross used to generate DAT-Flp;RCE:FRT mice. F-G) Coronal midbrain section showing DAT-Flp-dependent recombination of a Flp-dependent EGFP reporter (green) in DAT-Flp;RCE:FRT mice, demonstrating nearly complete overlap with endogenous TH immunostaining (magenta). White arrowhead indicates lack of EGFP expression in the interpeduncular nucleus (IPN). Panel G shows zoomed-in images of the VTA and SNc from the boxed regions in F. H-J) Confocal images of DAT-Flp-dependent recombination outside of the midbrain in a DAT-Flp;RCE-FRT mouse. The caudal linear nucleus raphe and arcuate nucleus show the expected partial overlap between the EGFP Flp reporter (green) and TH immunostaining (magenta) (white arrowheads). The dorsal raphe shows distinct EGFP labeling immediately ventral to the ventricle. These regions also contain TH+/EGFP^-^ neurons that are likely non-dopaminergic (magenta arrowheads). K) Charts showing the percentage of EYFP-positive neurons that co-express TH in different midbrain regions in DAT-Flp;RCE-FRT mice. >96% of EGFP+ neurons express TH in the SNc and VTA. Averaged from two P56 mice, 1 male and 1 female. L-M) Confocal images of sections of the locus coeruleus and zona incerta from a DAT-Flp;RCE-FRT mouse showing nearly no EGFP expression in TH+ neurons in these regions.

### Characterization of DAT-Flp mice

To determine whether Flp recombinase was efficiently expressed in DA neurons, adult heterozygous DAT-Flp mice were injected with an AAV expressing a Flp-dependent EYFP reporter in the midbrain (Fig. 1C). We observed the expected labeling of midbrain DA neurons in the injected area and greater than 97% of EYFP positive neurons were TH positive (Fig. 1D). Following this confirmation, we crossed the DAT-Flp mice to a mouse line expressing Flp-dependent EGFP (RCE:FRT) (31) (Fig. 1E). This cross resulted in Flp-dependent EGFP expression in canonical DA neuron-containing areas including the VTA, SNc, caudate linear nuclear raphe (CLi), dorsal raphe (DR), and arcuate nucleus (Fig. 1F-J). As expected, the vast majority (97%) of neurons in the SNc and VTA were TH positive (Fig. 1K), which matches expression patterns in the widely used DAT-IRES-Cre mouse line (22,26). There was no expression, however, in regions that contain TH-positive, DAT-negative neurons (32) such as the interpeduncular nucleus (Fig. 1F, white arrow, IPN), the locus coeruleus (LC, Fig. 1L), and the zona incerta/A13 (Fig. 1M). Together, these data demonstrate successful generation of a novel transgenic mouse line with robust Flp-dependent recombination in DA neurons without detectable off-target expression.

### The effects of Flp recombinase insertion into the DAT 3’ UTR

It has been reported in DAT-IRES-Cre mice that knock-in of Cre recombinase affects endogenous DAT expression. In adult heterozygous animals there is reduced DAT protein expression (26), reduced DA re-uptake, and increased peak-evoked DA concentration as measured by fast-scan cyclic voltammetry (FCV) (33). To test whether endogenous DAT expression is altered in DAT-Flp mice, which use a P2A sequence instead of an IRES sequence to achieve bicistronic expression that doesn’t disrupt exon 15, we compared DAT protein expression, DA release and re-uptake, and gross motor behaviors between DAT-Flp heterozygotes and their wild-type (WT) littermates.

Consistent with the previously observed reduction in DAT protein expression in DAT-IRES-Cre mice, which was replicated here (Supplemental Fig. 1A,B), we found a significant decrease in DAT protein in tissue punches from the dorsal striatum of heterozygous DAT-Flp mice compared to WT littermate controls (Fig. 2A,B). This change was selective for DAT as there were no significant differences in TH or VMAT2 expression, which is the vesicular monoamine transporter responsible for loading DA into vesicles (Fig. 2A,B). To measure DAT function, we used FCV to compare electrically-evoked DA release and re-uptake from WT and heterozygous DAT-Flp mice in ex vivo striatal slices. We measured single pulse-evoked DA release and re-uptake at seven locations throughout the striatum and nucleus accumbens (Fig. 2C). We found an average 32% increase in peak-evoked DA concentration across all dorsal striatum sites (dStr, sites 1-5, Fig. 2D) and a 30% increase in the nucleus accumbens core (NAc, sites 6+7, Fig. 2E) in DAT-Flp heterozygous mice. Using region- and peak concentration-matched transients, we observed slower DA re-uptake in DAT-Flp heterozygous mice (Fig. 2F) consistent with their reduced DAT protein expression. An increase in peak-evoked DA concentration and a small decrease in DA re-uptake rate were also observed in DAT-IRES-Cre mice (Supplemental Fig. 1C-F). Notably, the overall peak-evoked DA levels in WT mice from the two mouse lines were nearly three-fold different which likely reflects the differences in genetic background. The DAT-IRES-Cre mice were on a pure C57BL/6J background, while the DAT-Flp mice were on a CD-1/C57BL/6J mixed background. Previous work has shown differences in the dopaminergic system depending on mouse background strain (34-36). Therefore, these results show that DA release and re-uptake dynamics are highly sensitive to both DAT expression level and mouse genetic background.

**Figure 2.**
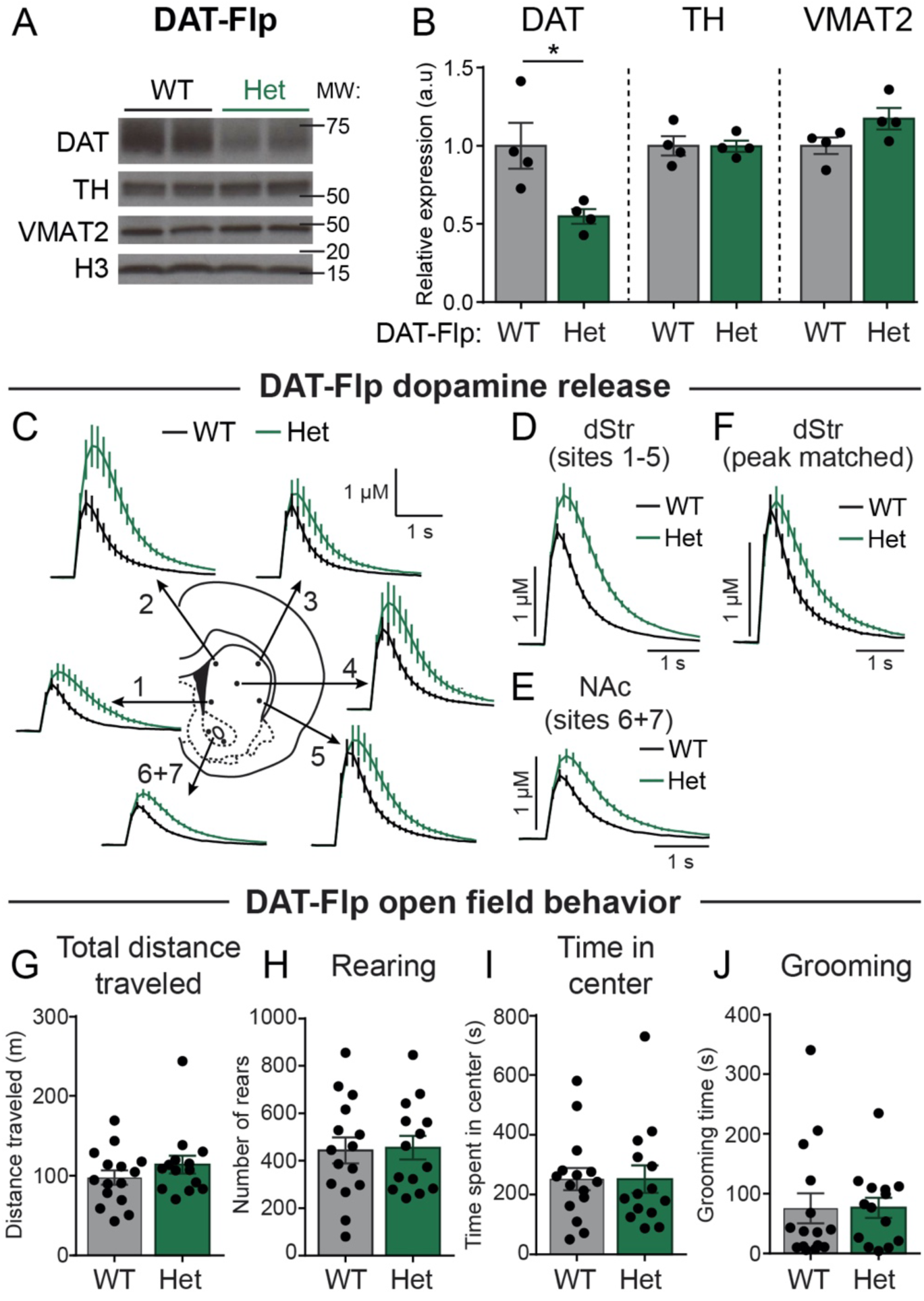
DAT expression and function in DAT-Flp mice. A) Representative western blot images of dopamine active transporter (DAT), tyrosine hydroxylase (TH), vesicular monoamine transporter 2 (VMAT2), and histone H3 (H3) loading control from striatal lysates from DAT-Flp wild-type (WT) and heterozygous (Het) mice. Two independent samples per genotype are shown. Blots were cropped to show the relevant bands. Molecular weight (MW) in KD is indicated on the right. B) Quantification of protein levels relative to histone H3, normalized to WT. Bars represent mean +/- SEM. Each dot represents the average of two striatal samples from one mouse (n=4 mice per genotype, 2 males and 2 females, P100-120). DAT: p=0.0259, TH: p=0.9639, VMAT2: p=0.0930; unpaired t-tests. C) Extracellular DA ([DA]_o_) transients evoked by single electrical pulses and recorded with fast-scan cyclic voltammetry (FCV) in different striatal sub-regions. Traces are mean +/- SEM [DA]_o_ versus time (average of 32 transients per recording site from 4 pairs of DAT-Flp WT and Het mice, 2 male pairs and 2 female pairs, P100-120). 1: ventromedial striatum, 2: dorsomedial striatum, 3: dorsolateral striatum, 4: central striatum, 5: ventrolateral striatum, 6-7: nucleus accumbens. D) Mean +/- SEM [DA]_o_ versus time (average of 80 transients per genotype) from the dorsal striatum sites #1-5. E) Mean +/- SEM [DA]_o_ versus time (average of 32 transients per genotype) from the nucleus accumbens sites #6-7. F) Region- and peak-matched mean +/- SEM [DA]_o_ versus time from the dorsal striatum of DAT-Flp WT and Het mice (average of 22 transients per genotype). DAT-Flp heterozygous mice have significantly slower [DA]_o_ re-uptake than WT mice (p<0.0001, single-phase exponential decay curve fit; WT tau=0.409, Het tau=0.648). G-J) Behavioral performance of DAT-Flp mice in a 60-minute open-field test. For all graphs, bars represent mean +/- SEM and dots represent values for individual mice. n= 15 WT mice, 6 males and 9 females; n=14 Heterozygous mice, 5 males and 9 females, P50-90. G) Total distance traveled in 60 minutes; p=0.2680, unpaired t-test. H) Total number of rears in 60 minutes; p=0.8847, unpaired t-test, I) Total time spent in the center of open field in 60 minutes; p=0.9981, unpaired t-test. J) Total number of grooming bouts in the first 20 minutes; p=0.8061, unpaired t-test. See also Supplemental Figure 1.

To test whether changes in DAT expression resulted in any gross behavioral alterations, we tested DAT-Flp mice of both sexes in the open field and measured locomotor activity, avoidance behavior, and repetitive behaviors (i.e. grooming). We found no significant difference in total distance traveled or number of rears between DAT-Flp heterozygous mice and WT littermate controls (Fig. 2G,H). We also found no changes in the time spent in center, a proxy for avoidance behavior, and no changes in time spent grooming for DAT-Flp mice (Fig. 2I,J). We tested DAT-IRES-Cre mice for comparison (Supplemental Fig. 1G-J) and did find a small but significant increase in locomotor activity and a decrease in grooming time in heterozygous mice compared to WT littermate controls. Together, these analyses show that even with use of a smaller self-cleaving P2A sequence instead of an IRES sequence, there can be changes in expression of the endogenous gene with a knock-in strategy. While this needs to be taken into consideration when designing experiments with DAT-IRES-Cre or DAT-Flp mice, our behavioral data show that changes in DAT do not strongly affect baseline motor activity or avoidance behavior. Therefore, DAT-Flp mice should be a useful tool for a variety of anatomical and behavioral experiments.

### Intersectional labeling of *Neurod6*-expressing midbrain DA neurons

Previous studies have revealed that DA neurons are composed of genetically distinct subclasses (12-15,37). One of these subpopulations, marked by the expression of the transcription factor *Neurod6*, is found in the ventromedial VTA and projects to the NAc medial shell (NAc MSh) (12,37-39). This subpopulation is of interest because it is selectively spared in a 6-OHDA model of Parkinson’s disease (12). Moreover, recent studies have identified distinct behavioral roles for VTA DA neurons that project to the MSh versus the lateral shell (LSh) of the NAc (8). In particular, LSh-projecting DA neurons show canonical dopaminergic responses to rewards and reward predictive cues, whereas MSh-projecting DA neurons respond more to aversive stimuli and aversive cues (8). Since both the cell bodies and projection sites of these two functionally distinct subpopulations are in close spatial proximity, it has been a challenge to selectively target them for further study. The restricted projections of *Neurod6*-expressing DA neurons, as previously defined by retrograde labeling and fluorescent in situ hybridization (FISH), make *Neurod6* a compelling marker gene to label MSh-projecting DA neurons (12,38). However, *Neurod6* is also highly expressed in excitatory neurons of the cortex, in nuclei adjacent to the VTA, and in non-dopaminergic neurons within the VTA (Fig. 3A-C) (12,28,40). Therefore, intersectional tools are needed to selectively and reliably target *Neurod6*-positive DA neurons.

**Figure 3.**
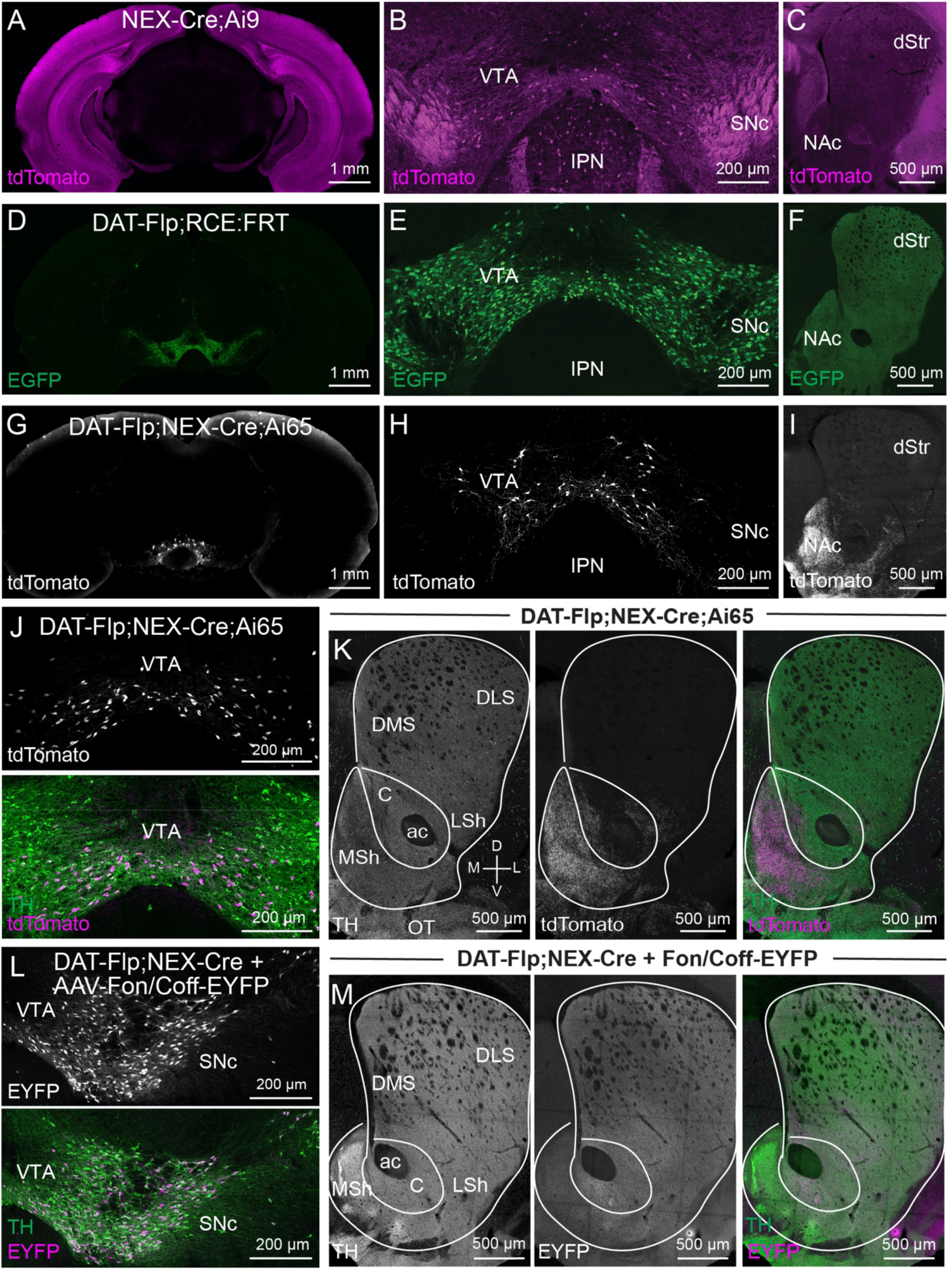
Intersectional genetic targeting of *Neurod6*-positive DA neurons. A-C) Confocal images from a P28 male NEX-Cre;Ai9 mouse showing tdTomato expression. A) Whole coronal section at A/P −3.4 from Bregma, showing high expression throughout the cortex and hippocampus. B) TdTomato expression in midbrain including the substantia nigra pars compacta (SNc) and ventral tegmental area (VTA). Note the fiber tracts in the medial lemniscus and cell bodies in the VTA and adjacent interpeduncular nucleus (IPN). C) TdTomato expression in the dorsal striatum (dStr) and nucleus accumbens (NAc) at A/P +1.34 from Bregma. D-F) Confocal images from a P50 male DAT-Flp;RCE:FRT mouse showing EGFP expression. D) Whole coronal section at A/P −3.4 from Bregma, showing restricted expression in the midbrain. E) TdTomato expression in the midbrain. F) TdTomato expression in the striatum and NAc at A/P +1.34 from Bregma. G-I) Confocal images from a P50 female DAT-Flp;NEX-Cre;Ai65 mouse showing tdTomato expression. G) Whole coronal section at A/P −3.4 from Bregma, showing restricted expression in the VTA. H) TdTomato expression in the midbrain. I) TdTomato expression in the striatum and NAc at A/P +1.34 from Bregma. J) Confocal images of a VTA section from a P50 DAT-Flp;NEX-Cre;Ai65 mouse showing tdTomato (upper panel) co-localization with TH immunostaining (lower panel, merged view). K) Confocal images of the projection targets of DAT-Flp;NEX-Cre;Ai65 neurons in the striatum and NAc. Left panel shows TH immunostaining. Middle panel shows tdTomato+ projections from DAT-Flp;NEX-Cre;Ai65 neurons. Right panel shows merged image with TH in green and DAT-Flp;NEX-Cre;Ai65 projections in magenta. [DMS = dorsomedial striatum, DLS = dorsolateral striatum, C = nucleus accumbens core, ac = anterior commissure, LSh = nucleus accumbens lateral shell, MSh = nucleus accumbens medial shell, OT = olfactory tubercle] L) Confocal images of the SNc/VTA of a P90 female DAT-Flp;NEX-Cre mouse injected with AAV-Fon/Coff-EYFP (top panel) with TH immunostaining (bottom panel, merged view). M) Confocal images of the projection targets of DAT-Flp;NEX-Cre neurons expressing Fon/Coff-EYFP in the striatum and NAc. Left panel shows TH immunostaining. Middle panel shows EYFP+ projections from DAT-Flp-positive/NEX-Cre negative neurons. Right panel shows merged image with TH in green and DAT-Flp-positive/NEX-Cre-negative projections in magenta. See also Supplemental Figure 2.

To access *Neurod6*-positive DA neurons, we crossed DAT-Flp;Ai65 mice (Fig. 3D-F) to NEX-Cre mice (28), which label the majority of *Neurod6*+ neurons (NEX is the former name for *Neurod6*). In the resulting triple transgenic offspring (Supplemental Fig. 2A), we found remarkably specific tdTomato labeling of a small population of neurons in the ventromedial VTA (Fig. 3G,H,J). The anatomical location of these Neurod6*/*DAT*+* neurons matched closely to what was previously observed by FISH (12). As predicted by retrograde tracing (12), we found that *Neurod6*/DAT+ neurons projected strongly to the NAc MSh, with some additional projections going to the olfactory tubercle (OT) and lateral septum (Fig. 3I,K, Supplemental Fig. 2B). We confirmed that the Neurod6/DAT+ neurons and projections from these mice expressed TH (95% co-expression, Fig. 3J,K).

To test whether dopaminergic projections to the NAc MSh arise exclusively from the *Neurod6+* subpopulation, we injected an AAV expressing a Flp-on/Cre-off-EYFP construct (20) into the midbrain of DAT-Flp;NEX-Cre mice (Supplemental Fig. 2C). In this experiment, only DA neurons that do not express NEX-Cre will be labeled with EYFP (Fig. 3L). We observed the expected EYFP axonal arborizations throughout the dorsal striatum, with notably reduced projections in the NAc MSh and OT (Fig. 3M). This analysis demonstrates that *Neurod6*+ DA neurons make up a large majority of NAc MSh-projecting DA neurons. Together, these data highlight the utility of viral and transgenic intersectional strategies to label subpopulations of DA neurons based on their expression of Flp and Cre recombinase.

### Whole-brain imaging of *Neurod6+* DA neurons

To measure the projection patterns of *Neurod6*+ DA neurons in an unbiased, brain-wide manner, we optically cleared whole brains from DAT-Flp;NEX-Cre;Ai65 mice using the iDISCO - based (41) brain clearing method Adipo-Clear (29) and imaged the intact brain using a light sheet microscope (Fig. 4 and Supplemental Video 1). We confirmed the remarkable specificity of the intersectional strategy, with tdTomato+ cell bodies primarily located in the VTA with clearly defined projections to the NAc MSh (Fig. 4A-E). We further analyzed the tdTomato+ projections of *Neurod6+* DA neurons using TrailMap (42). The NAc MSh projections were too dense and complex to computationally recognize as individual fibers (Fig. 4D) and were excluded from the quantification. The most numerous projections outside of the NAc were found in the OT, with additional sparse projections in the hypothalamic lateral zone (lateral preoptic area, LPO; preparasubthalamic nucleus, PST; parasubthalamic nucleus, PSTN; perifornical nucleus, PeF; Supplemental Fig. 3A, Supplemental Table 1 and Supplemental Videos 2 and 3). TdTomato+ processes were also detected in the midbrain (paranigral nucleus, PN; retrorubral area, RR) and the raphe nuclei (interpeduncular nucleus, IPN; rostral linear nucleus raphe, RL; central linear nucleus raphe, CLi), which likely represent the dendritic arbors of *Neurod6+* DA neurons (Supplemental Fig. 3A). Together, these data show that the vast majority of projections from *Neurod6*+ DA neurons go to the NAc MSh with minor projections to the OT and hypothalamus. These projection patterns are reproducible and consistent across multiple mice (Fig. 4D,E and Supplemental Fig. 3A).

**Figure 4.**
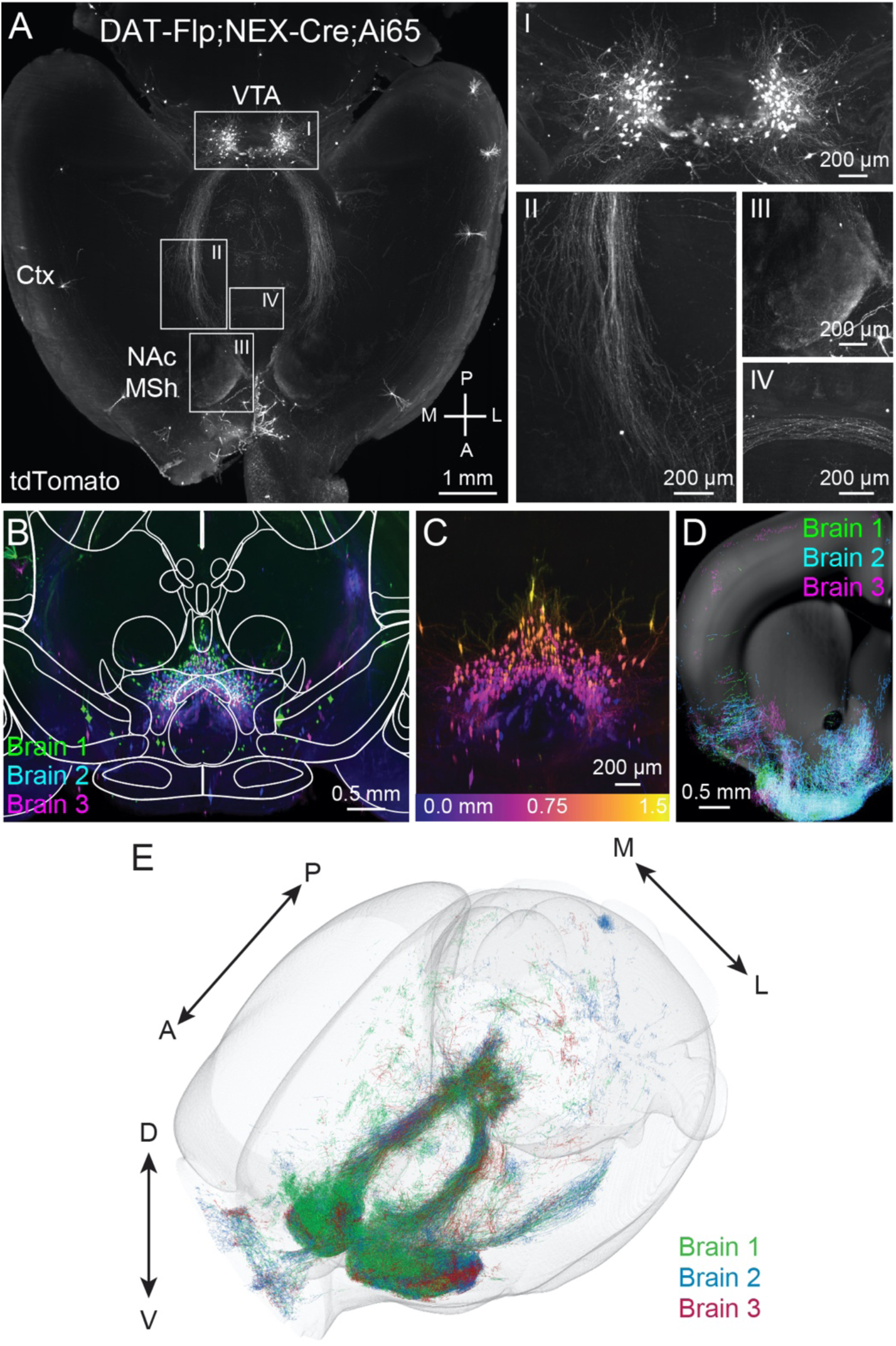
Whole brain imaging of DAT-Flp;NEX-Cre;Ai65 mice. A) Whole brains from 2 male and 1 female P120 DAT-Flp;NEX-Cre;Ai65 mice were optically cleared and processed using the Adipo-Clear pipeline. TdTomato fluorescence was amplified with an anti-RFP antibody and whole brains were imaged without sectioning using a light-sheet microscope. Shown is a max projection of 600 μm from a horizontal plane Z-stack. Right panels show zoomed-in images of *Neurod6*+ DA neurons in the midbrain (I), axonal tracts from the midbrain to the nucleus accumbens medial shell (NAc MSh) through the medial forebrain bundle (II), *Neurod6*+ DA neuron axon terminals in the MSh (III), and contralateral fibers crossing the midline (IV). Ctx = cortex. B) Coronal view (XZ projection) comprising a 1.5 mm A/P cross section of the VTA in optically cleared DAT-Flp;NEX-Cre;Ai65 mice. 3 separate brains were aligned and merged. Image is overlaid with brain region outlines from the middle position of the 1.5 mm cross-section from the Allen Brain Institute reference atlas. C) Coronal Z-stack image of a 1.5 mm cross section of the VTA from a single brain color coded by depth. Depth scale is shown in lower panel. D) Coronal Z-stack image of TrailMap-extracted axons from a 500 μm section of the striatum and NAc. 3 separate brains were aligned and merged. Extracted axons are overlaid onto a reference slice from the middle position of the 500 μm section from the Allen Brain Atlas. E) Projections and processes of DAT-Flp;NEX-Cre;Ai65-positive neurons visualized in a 3D view of TrailMap-extracted processes from three aligned brains. See also Supplemental Figure 3, Supplemental Table 1, and Supplemental Videos 1-3

We did observe a small number of tdTomato^+^ neurons outside of traditional dopaminergic regions including in the cortex and hippocampus (Fig. 4A). To test whether these neurons were bona fide *Neurod6+* DA neurons or reflected off-target recombination events, we performed antibody staining against TH. None of the cortical neurons were TH positive (Supplemental Fig. 3B). Given the robust expression of *Neurod6* throughout the cortex (see Fig. 3A) and lack of EGFP positive cortical neurons in DAT-Flp;RCE-FRT mice (see Fig. 3D), these neurons may represent NEX-Cre-positive neurons with leaky recombination events. Alternatively, they may reflect transient cortical DAT-Flp expression leading to recombination of the Ai65 tdTomato reporter. In addition to the cortex, we also found small populations of tdTomato+ cells in the anterior olfactory area and cerebellum (Supplemental Fig. 3C,D and Supplemental Video 1).

### Functional specificity of DA release from *Neurod6*+ DA neurons

The anatomical data described above indicate a robust and selective projection from *Neurod6+* DA neurons to the NAc MSh. To confirm the functionality of this projection, we bilaterally injected an AAV expressing Flp- and Cre-dependent ChR2 (20) into the midbrain of DAT-Flp;NEX-Cre mice (Fig. 5A-C). To assess whether activation of *Neurod6+* neurons leads to DA release, we measured DA levels with FCV in response to optical stimulation of ChR2+ terminals in different subregions of the striatum. We observed robust optically-evoked DA release in the NAc MSh (Fig. 5D). MSh DA release was reliable as we observed stable peak-evoked DA levels with repeated stimuli over the course of 20 minutes (Fig. 5D). Optical stimulation outside the MSh did not result in measurable DA transients, consistent with the lack of ChR2 expression in those regions (Fig. 5C,D). These results confirm the utility of DAT-Flp mice for future functional assays using optogenetics and demonstrate the specificity of intersectional approaches to label discrete dopaminergic subpopulations.

**Figure 5.**
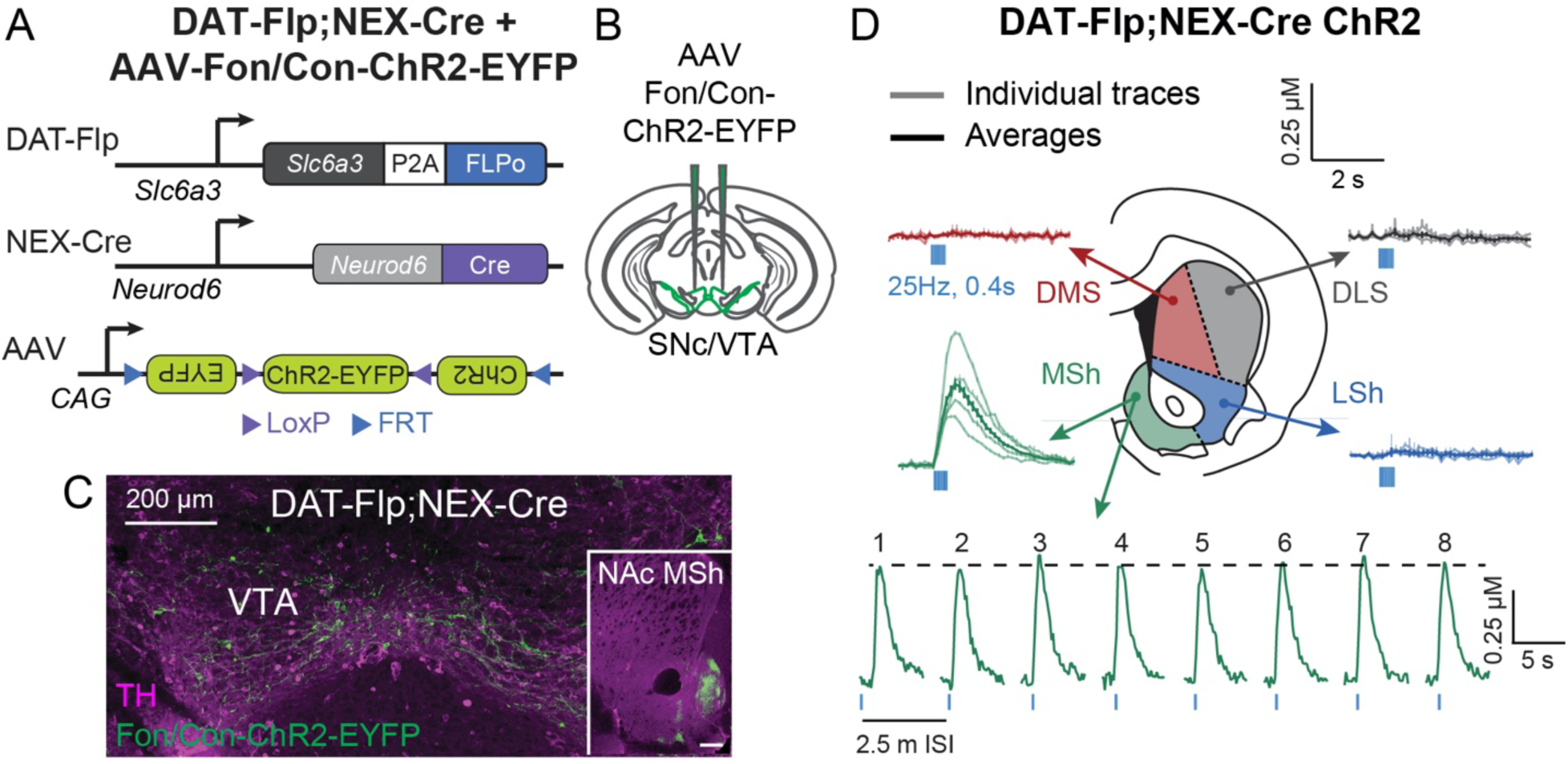
Optogenetic activation of DA release in DAT-Flp;NEX-Cre mice. A) Schematic of the intersectional genetic strategy to target *Neurod6*+ DA neurons in DAT-Flp;NEX-Cre mice with an AAV expressing Flp- and Cre-dependent channelrhodopsin (ChR2). B) Schematic showing bilateral injection of AAV-Fon/Con-ChR2-EYFP into the midbrain of DAT-Flp;NEX-Cre mice. SNc = substantia nigra pars compacta; VTA = ventral tegmental area. C) Confocal image showing midbrain expression of Fon/Con-ChR2-EYFP (green) and co-expression with TH immunostaining (magenta) in a DAT-Flp;NEX-Cre mouse. Inset shows ChR2-EYFP+ projections in the nucleus accumbens medial shell (NAc MSh) in an injected mouse. Scale bar represents 200 μm. D) Fast-scan cyclic voltammetry traces of optically-evoked extracellular DA ([DA]_o_) in striatal slices from DAT-Flp;NEX-Cre mice injected with Fon/Con-ChR2-EYFP. Mean +/- SEM [DA]_o_ versus time evoked from different striatal subregions by 10 light pulses delivered at 25 Hz. Light colored lines show individual traces (n=3-9 transients per recording site) and dark colored lines are the average of all traces, recorded from 4 hemispheres of 2 P120 male mice. DMS = dorsomedial striatum, DLS = dorsolateral striatum, LSh = nucleus accumbens lateral shell, MSh = nucleus accumbens medial shell. Bottom panel shows transients recorded in the MSh from 8 consecutive stimulations delivered 2.5 minutes apart.

## Discussion

Here we describe a novel transgenic mouse line expressing Flp recombinase from the endogenous DAT locus, which provides a new tool for the visualization and manipulation of DA neurons. We provide anatomical validation of this new mouse line by demonstrating robust and specific Flp-mediated recombination of genetic and viral reporters in TH-positive DA neurons. We further show functional validation that Flp-dependent ChR2 can be used to evoke DA release. We demonstrate proof-of-concept that DAT-Flp mice can be crossed to mice expressing Cre under specific gene promoters to allow for the intersectional targeting of dopaminergic subpopulations defined by their co-expression of DAT and any other gene of interest.

Single-cell sequencing studies of neuronal populations throughout the brain have provided unprecedented access to the gene expression profiles of different cell types. These studies have highlighted the need for genetically targeted approaches to enable the study of neuronal subpopulations in a specific and reproducible way (43). This is especially important for heterogeneous cell types in which spatially adjacent or intermingled neurons can have different functional roles, thus constituting distinct subtypes, such as in the striatum or midbrain dopaminergic system (8,9,44-48). In recent years, there have been major efforts to generate mouse lines that express Cre recombinase from different cell type-specific promoters (49-51). These mouse lines have become essential tools for neuroscience and other areas of biology. However, given the limited repertoire of genes that are expressed in neurons, there is often substantial expression of ‘marker’ genes in more than one cell type. Therefore, single genes often aren’t sufficient to label specific neuronal subtypes within functionally diverse neuronal populations.

Mouse genetic intersectional strategies, based on the expression of different recombinases under the control of two or more genes, represent emerging approaches that greatly increase our ability to target newly defined neuronal subpopulations (20,21). This strategy has been successfully applied to neuromodulatory populations, including serotonergic and dopaminergic neurons (18,19). The DAT-Flp mouse line described here contributes to this growing toolkit of intersectional mouse genetic resources. For labeling DA neurons specifically, DAT-Flp mice have some advantages over currently available intersectional mouse models, such as the Th-2A-Flpo mouse (18). Namely, DAT drives Flp expression selectively in DA neurons, whereas Th-2A-Flpo is expressed in catecholaminergic neurons throughout the brain. This an important consideration when attempting to target midbrain DA neurons with viral or genetic approaches as TH is expressed in non-dopaminergic neurons in the interpeduncular nucleus and the linear raphe, which reside just adjacent to the VTA (22,32). In addition, TH-positive neurons are found in the locus coeruleus, dorsal raphe, hypothalamus, and zona incerta, among others (32). Therefore, Th-2A-Flpo mice label a larger population of neurons. Together DAT-Flp and Th-2A-Flpo mice represent complementary tools that will have different utility depending on which neuronal population is to be targeted.

Here we find that similar to the commonly used DAT-IRES-Cre mouse line (26), bicistronic expression of Flp from the endogenous *Slc6a3* locus alters DAT protein expression. We found a reduction in DAT levels in the dorsal striatum in mice that were heterozygous for DAT-Flp, which resulted in detectable changes in peak evoked DA release and re-uptake kinetics in the striatum. While the mechanism for this is unknown, it may result from loss of the 3’ UTR or the instability of the long mRNA transcript that is produced from the targeted locus, which contains the coding sequence for both DAT and Flp. This change in DAT expression will need to be considered when designing experiments with DAT-Flp mice, especially if measures related to DAT function are being explored. That said, we did not find any changes in gross locomotor activity in DAT-Flp heterozygous mice, suggesting that they should be useful for many types of experiments. We suggest designing experiments so that all mice are DAT-Flp heterozygous, and thus have similar DAT expression, and avoid comparing DAT-Flp heterozygotes directly with WT mice.

In addition to characterizing the DAT-Flp mice, we highlight an example of how an intersectional genetic approach can be used to visualize and manipulate a specific subpopulation of DA neurons. This population is marked by the expression of a transcription factor *Neurod6*, which is expressed in about 25% of DA neurons in the ventromedial VTA and was previously shown to comprise the majority of projections to the NAc MSh by retrograde labeling (12). This population is of considerable interest given recent studies showing that *Neurod6* marks VTA DA neurons that are spared in a 6-OHDA model of Parkinson’s disease (12), and could be neuroprotective by maintaining mitochondrial health (52,53). In addition, *Neurod6* and its related family member *Neurod1* have been shown to be important for the developmental survival of this VTA subpopulation (39).

At the behavioral level, there is strong evidence that MSh-projecting DA neurons in the VTA are activated by unexpected aversive events (8). However, it has also been reported that high power optogenetic activation of NEX-Cre-expressing neurons in the midbrain can lead to real-time place preference, signifying a rewarding outcome (38). Moreover, it has been shown that while increased DA release in the NAc core may signal reward, this is not necessarily correlated with cell body firing in the VTA (54). Axonal DA release can be gated by local mechanisms independent of cell body firing (55); however, without stable and consistent labeling of the same DA population it is difficult to confirm the relationship between the activity of VTA neurons and axonal DA release at the projection target. These variable and somewhat conflicting results highlight the need for consistent and tractable marking of dopaminergic subpopulations without the labeling of off-target cell types. The DAT-Flp;NEX-Cre mice described here represent a new model that will allow selective access to this behaviorally and disease-relevant subpopulation of DA neurons.

## Materials and Methods

### Mice

Animal procedures were conducted in accordance with protocols approved by the University of California, Berkeley Institutional Animal Care and Use Committee (IACUC) and Office of Laboratory Animal Care (OLAC).

The DAT-Flp mice were generated at UC Berkeley as described below. DAT^IRES^Cre mice (Jackson Laboratory, strain #006660 (26)) were crossed and maintained with C57BL/6J wild-type mice (Jackson Laboratory, strain #000664). RCE:FRT mice (31) were used to endogenously label Flp-expressing neurons with EGFP (Gt(ROSA)26Sortm1.2(CAG-EGFP)Fsh/Mmjax, MMRRC stock #32038-JAX). To label neurons co-expressing Flp and Cre recombinase, the Ai65 mouse line (30) was used (Ai65(RCFL-tdT)-D, Jackson Laboratory strain #021875). To target *Neurod6* expressing neurons, we used the NEX-Cre mouse line (NEX was the previous name of *Neurod6*) obtained from Dr. Klaus-Armin Nave (28). Mice were housed on a 12 h light/dark cycle and given ad libitum access to standard rodent chow and water. The ages, sexes, and numbers of mice used are indicated for each experiment in figure legends.

### Generation of DAT-Flp mice

DAT-Flp mice were generated from Ai65 mouse embryonic stem cells (mESCs) from the Allen Institute for Brain Science (a gift from Tanya Daigle, Ph.D.). Ai65 mESCs harbor a Flp- and Cre-dependent tdTomato fluorescent reporter in the ROSAβgeo26 locus (30). Ai65 mESCs were karyotyped (Cell Line Genetics, Inc. Madison, WI) to ensure chromosomal health both before and after gene editing. Clonal cell lines that showed normal chromosome numbers and no deletions/insertions in at least 19 of 20 cells analyzed were considered to be chromosomally normal.

To generate the in-frame genomic insertion of a P2A-Flpo sequence, we used CRISPR-Cas9 to generate a double-stranded break immediately upstream of the stop codon in the last exon (exon 15) of the *Slc6a3* gene. To do this, we designed two plasmids: a donor plasmid and a Cas9 plasmid. For the donor construct, we generated a 2499 bp gBlock gene fragment (Integrated DNA technologies, IDT, sequence below) that contained a 15 bp overhang for integration into a plasmid, a 552 bp homology arm on the 5’ end, a P2A sequence starting with a GSG (56), a nuclear localization sequence, the optimized Flp-recombinase (Flpo) sequence (30,57), a 552 bp homology arm on the 3’ end, and a 15 bp overhang for integration into a plasmid. This sequence was inserted into a donor plasmid (pCRII-TOPO) using In-Fusion HD cloning (Takara – 639649). For the Cas9 plasmid, a single-guide RNA (sgRNA) targeting the 5’ NGG protospacer adjacent motif (PAM) immediately downstream of exon 15 (sgRNA forward: 5’ *CACCG*CATTGGCTGTTGGTGTAAG 3’, sgRNA reverse: 5’ *AAAC*CTTTACACCAACAGCCAATG*C* 3’, *italic* portions represent sequences present for px330 plasmid insertion) was inserted into the CRISPR-Cas9-encoding px330 plasmid (58).

The px330 plasmid along with the donor constructs were nucleofected into 5×10^6^ Ai65 mESCs. Transfected cells were cultured and the resulting colonies were screened for correct insertion and integration. We designed a genotyping strategy with a forward primer immediately upstream of the first homology arm in exon 15 of *Slc6a3* (DAT-Flp geno forward: 5’ CATGCAGAAGGACAGACACT 3’), a reverse primer in the Flp sequence 650 bp away from the forward primer (Flp insert reverse: 5’ AGGATGTCGAACTGGCTCAT 3’), and a reverse primer in the 3’ UTR of the *Slc6a3* gene, 1,100 bp away from the forward primer when there is no insert (DAT-Flp geno reverse: 5’ ACCCTGCGTGTGTGTAATAT 3’). Flp insertion was confirmed by the presence of a 650 bp PCR product (Fig. 1B). With this strategy, the forward primer falls outside of the homology arm, therefore we could confirm on-target insertion.

Following confirmation of correctly targeted Ai65 mESC clones with normal karyotypes, mESCs were given to the Berkeley Cancer Research Laboratory Gene Targeting Facility for the generation of chimeras using aggregation of mESCs and morula embryos as described previously (59). Pups born from these females were checked for chimerism by coat color and tail DNA genotyping. F1 pups and subsequent generations were genotyped using tail DNA.

DAT-P2A-FlpO gBlock gene fragment:

<u>TACCGAGCTCGGATC</u>GGACCACTCACAAATCAGTAGGAGGACAATATGGGGCCAGTCCTGCATCTAAAGAT GTTCTGCATATCTCACCTCAGCTACTGATCACAAATATGCTGGGCAGAAAGAACCCTGGCCTTCTAAGCACCTGGGCTCCATGAGGCTTGGTTATTTGTTGGTTCTGGTTTGTTTTCCTTTACTCTGCCTTTTTTTTTGTTTGTTTGTTTGTTTGTTTTGAGACAATCTTACTGTAGCACAGGCTGGCCTCAAACCCATACCTGCTAGCCTCTTGAGATCATGACTACAGACATTAGCTGCTCTACCAAGTTTACAGGTCCCTTCTTCATCAAGTAATTCAAGTACTTAAAAGTTTCCATGAGGTCATTAGCCGTTGGTGCCCTCAAGGGTGGTATCTATCATCGGTAACCTCCCCCATGGGCAAGTAGTACCACCTGCTGAATTGATCCTGATTTCATCAGCGAGTCAGTGGCCCATAAGCCGTGATTCTAAAGTCTGCTTCTCACTCGTAAATAAATGATGCTTTCTTTCCAGCTGCGCCATTGGCTGTTGGTGGGAAGCGGAGCTACTAACTTCAGCCTGCTGAAGCAGGCTGGAGACGTGGAGGAGAACCCTGGACCTATGGCTCCTAAGAAGAAGAGGAAGGTGATGAGCCAGTTCGACATCCTGTGCAAGACCCCGCCGAAGGTGCTGGTGCGGCAGTTCGTGGAGAGATTCGAGAGGCCCAGCGGCGAAAAGATCGCCAGCTGTGCCGCCGAGCTGACCTACCTGTGCTGGATGATCACCCACAACGGCACCGCGATCAAGAGGGCCACCTTCATGAGTTATAACACCATCATCAGCAACAGCCTGAGTTTTGACATCGTGAACAAGAGCCTGCAGTTCAAGTACAAGACCCAGAAGGCCACCATCCTGGAGGCCAGCCTGAAGAAGCTGATCCCCGCATGGGAGTTCACGATTATCCCTTACAACGGCCAGAAGCACCAGAGCGACATCACCGACATCGTGTCCAGCCTGCAGCTGCAGTTCGAAAGCAGCGAGGAGGCCGACAAGGGGAATAGCCACAGCAAGAAGATGCTGAAGGCCCTGCTGTCCGAAGGCGAGAGCATCTGGGAGATTACCGAGAAGATCCTGAACAGCTTCGAGTACACCAGCAGATTTACCAAAACGAAGACCCTGTACCAGTTCCTGTTCCTGGCCACATTCATCAACTGCGGCAGGTTCAGCGACATCAAGAACGTGGACCCGAAGAGCTTCAAGCT CGTCCAGAACAAGTATCTGGGCGTGATCATTCAGTGCCTGGTCACGGAGACCAAGACAAGCGTGTCCAGGCACATCTACTTTTTCAGCGCCAGAGGCAGGATCGACCCCCTGGTGTACCTGGACGAGTTCCTGAGGAACAGCGAGCCCGTGCTGAAGAGAGTGAACAGGACCGGCAACAGCAGCAGCAACAAGCAGGAGTACCAGCTGCTGAAGGACAACCTGGTGCGCAGCTACAACAAGGCCCTGAAGAAGAACGCCCCCTACCCCATCTTCGCTATTAAAAACGGCCCTAAGAGCCACATCGGCAGGCACCTGATGACCAGCTTTCTGAGCATGAAGGGCCTGACCGAGCTGACAAACGTGGTGGGCAACTGGAGCGACAAGAGGGCCTCCGCCGTGGCCAGGACCACCTACACCCACCAGATCACCGCCATCCCCGACCACTACTTCGCCCTGGTGTCCAGGTACTACGCCTACGACCCCATCAGTAAGGAGATGATCGCCCTGAAGGACGAGACCAACCCCATCGAGGAGTGGCAGCACATCGAGCAGCTGAAGGGCAGCGCCGAGGGCAGCATCAGATACCCCGCCTGGAACGGCATTATAAGCCAGGAGGTGCTGGACTACCTGAGCAGCTACATCAACAGGCGGATCTGAAGTGGAAGGAGACAGCTGCCAGCTGGGCCATCTCACAACAGCGGGGACAGGGAGATCACAAAGGAAACCAACACGTCAAGAAAGGAGGGCCACTTCCACAGTCCCCTTTTGCCATATGGAAAAATAATCCAAGCATGGGCTTCAATCTTTGACTGTTCACACCCAATCCATGCCACAAAGAAGCCTCTGTCTGTGTGTGACTGTAAAAACACACACCTCTATACAGTGAAGTCAACCATGTCCCTGTCCCTAATGGGTGGGGAAACCCCTAGCTGGTATCCTGTCCTGCAAGGCTGACTCCCCCATCTGTGGTCACTCTGGGAGAACAGGTCATACTGTTCCCTGCATTCTAGGAGAGGGACTTTGGTACCTGTATATACACTGTGCCAGAATCCTGTGCTCACGGTAGTTGCCTAGATAATTTCTTTTGCTTAAATTTACAGTGTCAAGTATCCTATTTTTTGCTGTTGGTAGAAAAGACAGTTAATACATGCCAAGTCCTTTCCTGGTGCTTGGCTCCGAGCAGACACCATGACCTT AGCATCCTGTTCA<u>TCGAGCATGCATCTA</u>

### Stereotaxic intracranial injections

Intracranial injections were performed on male and female postnatal day (P) 40-60 mice. Mice were briefly anesthetized with 3% isoflurane (Piramal Healthcare – PIR001710) and oxygen. Their heads were shaved and mice were mounted on a stereotaxic frame (Kopf instruments – Model 940) with stabilizing ear cups and a nose cone delivering constant 1.5% isoflurane in medical oxygen. Viruses were injected using a pulled glass capillary at a rate of 100 nl/minute. Following injection, the capillary remained in place for 1 minute per every minute spent injecting to allow the tissue to recover and prevent virus backflow up the injection tract upon retraction. The following coordinates from Bregma were used to target each site: SNc (M/L ±1.5 mm, A/P - 2.8 mm, D/V −4.0 mm), VTA (M/L ±0.6 mm, A/P −3.1 to −3.7 mm for multiple injections, D/V −4.2 mm).

For ChR2 experiments, 400 nl of AAV8 Flp- and Cre-dependent ChR2 (pAAV-hSyn Con/Fon hChR2(H134R)-EYFP: Addgene – 55645-AAV8: titer 2.1 × 10^13^) diluted 1:7 in sterile saline was injected bilaterally into the anterior and posterior VTA of DAT-FLP;Nex-Cre mice. For DAT-Flp EYFP labeling experiments, 400 nl of undiluted AAV5 Flp-dependent EYFP (rAAV5.hSyn-Coff/Fon-eYFP-WPRE: UNC vector core: titer 3.5 × 10^12^) was injected bilaterally into the SNc and VTA of DAT-Flp heterozygous mice. For Intersectional labeling of *Neurod6*-negative DA neurons, 400 nl of AAV5 Coff/Fon-EYFP (rAAV5.hSyn-Coff/Fon-eYFP-WPRE: UNC vector core: titer 1.7 × 10^12^) diluted 1:1 in sterile saline was injected bilaterally into the SNc and the VTA of DAT-Flp;Nex-Cre mice.

### Brain sectioning and immunohistochemistry

Adult mice were anesthetized with isoflurane and transcardial perfusion was performed with 7.5 ml of 1x PBS following by 7.5 ml of ice cold 4% PFA in 1x PBS (EMS – 15710-S). Brains were then dissected out and post-fixed in 4% PFA in 1x PBS for 2 hours at 4°C. Brains were transferred to cryoprotectant solution (30% sucrose in 0.1M PB) at 4°C until they sank. 30 μm coronal sections were made using a freezing microtome (American Optical, AO 860). Sections were collected in serial wells of a 24 well plate in 1x PBS with 0.02% sodium azide (NaN_3_; Sigma-Aldrich – 26628-22-8) and stored at 4°C. For immunohistochemistry, individual wells of sections were washed with gentle shaking for 3 × 5 minutes with 1x PBS, then blocked for 1 hour at RT with BlockAid blocking solution (Life Tech: B10710). Primary antibodies diluted in PBS-Tx (1x PBS with 0.25% triton-X-100 (Sigma – T8787)) were then added and the tissue was incubated for 48 h with gentle shaking at 4°C. Sections were then washed with gentle shaking 3 × 10 minutes with PBS-Tx. Secondary antibodies diluted in PBS-Tx were added and incubated with shaking for 1 h at room temperature. Sections were then washed 3 × 10 minutes gently shaking with 1x PBS. Sections were mounted onto SuperFrost slides (VWR: 48311-703) and coverslipped with hard-set Vectashield mounting media (Vector Labs: H-1500). The following antibodies were used: anti-tyrosine hydroxylase (TH, 1:1000, mouse, Immunostar #22941), anti-RFP (1:1000, rabbit, Rockland #600-401-379), Alexa Fluor 488 goat anti-mouse secondary (1:500, Thermo Fisher Scientific #A-11001), Alexa Fluor 633 goat anti-mouse secondary (1:500, Thermo Fisher Scientific #A-21050), and Alexa Fluor 546 goat anti-rabbit secondary (1:500, Thermo Fisher Scientific #A-11035).

### Confocal microscopy and image analysis

Images of 30 μm sections processed for immunohistochemistry were acquired using an Olympus FV3000 confocal microscope equipped with 405, 488, 561, and 633 nm lasers and a motorized stage for tile imaging. Z-stack images captured the entire thickness of the section at 2-2.5 μm steps for images taken with a 20x air objective (Olympus #UCPLFLN20X) and 10 μm steps for images taken with a 4x air objective (Olympus #UPLXAPO4X).

Quantification of EGFP overlap with TH immunolabeling in DAT-Flp:RCE:FRT mice was done on max projected Z-stack images from 2 hemispheres per mouse and averaged together. Every clearly identifiable EGFP^+^ neuron in the SNc and VTA was manually counted using the Cell Counter Image J (NIH) plug-in. A neuron was considered TH^+^ if there was a clear TH signal in the entirety of the EGFP^+^ cell body. Boundaries of each sub-region of the VTA and SNc were made according to the Paxinos mouse brain atlas (60).

### Western Blotting

Bilateral punches (3 mm diameter) were taken from 275 μm thick striatal sections prepared from male and female (P50-90) DAT-Flp and DAT-IRES-Cre mice. Sections were cut on a vibratome in cold ACSF-Sucrose cutting solution (In mM: 85 NaCl, 25 NaHCO_3_, 2.5 KCl, 1.25 NaH_2_PO_4_, 0.5 CaCl_2_, 7 MgCl_2_, 10 glucose, and 65 sucrose). Punches contained dorsal striatum and small neighboring portions of the lateral septum and nucleus accumbens. Tissue punches were put into a 1.5 ml tube and frozen on dry ice. Samples were resuspended in 300 μl lysis buffer (1x PBS, 1% SDS, 1% Triton-X-100, 2 mM EDTA, and 2 mM EGTA) with Halt phosphatase buffer inhibitor (Fisher: PI78420) and Complete mini EDTA-free protease inhibitor (Roche: 4693159001). Samples were sonicated at low power (Qsonica Q55) until homogenized. Homogenized samples were then boiled for 5 minutes, spun down, and put on ice. Total protein concentration was determined using a BCA assay (Fisher: PI23227).

4x Laemmli sample buffer (Bio-Rad: 161-0747) with 5% 2-mercaptoethanol (Thermo Fisher Scientific) was added 3:1 to an aliquot of each sample. 15 μg of total protein were loaded onto a 4-15% Criterion TGX gel (Bio-Rad: 5671084). Proteins were transferred to an activated PVDF membrane (BioRad: 1620177) at 4°C overnight using the Bio-Rad Criterion Blotter. Membranes were then removed, cut, washed, and blocked in 5% milk in 1x TBS + 1% Tween-20 (TBS-T) for 1 h at RT. Blots were incubated with primary antibodies diluted in TBS-T + 5% milk overnight at 4°C. The next day, the blots were washed 3 × 10 minutes with TBS-T followed by 1 hour of incubation at RT with HRP-conjugated secondary antibodies (1:5000). Membranes were washed 6 x 10 minutes with TBS-T and incubated for one minute with a chemiluminesence substrate (Perkin-Elmer: NEL105001EA) and exposed to GE Amersham Hyperfilm ECL (VWR: 95017-661). Membranes were then stripped 2 x 6 minutes with 6 mM Guanidinium Chloride + 5% 2-mercaptoethanol, washed with TBS-T, and incubated with primary antibody overnight at 4°C as before.

The following primary antibodies were used for western blotting: mouse anti-tyrosine hydroxylase (TH, 1:3000, Immunostar - 22941); mouse anti-Histone-3 (1:1500, Cell Signaling - 96C10); rabbit anti-VMAT2 (1:1400, Alomone Labs - AMT-006), mouse anti-Dopamine Transporter [6V-23-23] (DAT, 1:1,400, Abcam – 128848). The following secondary antibodies were used: goat anti-rabbit HRP (1:5000, Bio-Rad: 170-5046) and goat anti-mouse HRP (1:5000, Bio-Rad: 170-5047)

### Fast-Scan Cyclic Voltammetry

DA release was monitored using fast-scan cyclic voltammetry (FCV) in acute coronal slices. Paired site-sampling recordings were performed in one WT reference animal and one heterozygous experimental animal from a sex- and age-matched mouse pair recorded on the same day. The genotype order of the tissue prep and recording chamber position were counterbalanced between experiments. Male and female mice (P50-90) were deeply anesthetized by isoflurane and decapitated. 275 μm thick coronal striatal slices were cut on a vibratome (Leica VT1000 S) in ice cold ACSF-Sucrose cutting solution (in mM: 85 NaCl, 25 NaHCO_3_, 2.5 KCl, 1.25 NaH_2_PO_4_, 0.5 CaCl_2_, 7 MgCl_2_, 10 glucose, and 65 sucrose). Slices recovered for 1 h at RT and were recorded from in ACSF (in mM: 130 NaCl, 25 NaHCO3, 2.5 KCl, 1.25 NaH2PO4, 2 CaCl2, 2 MgCl2, and 10 glucose). All solutions were continuously saturated with 95% O_2_ and 5% CO_2_. Slices between +1.2 mm and +0.5 mm (A/P) from Bregma that had both dorsal striatum and the entire nucleus accumbens were used for experimentation.

Prior to recording electrically- or light-evoked transients, ex vivo slices were equilibrated to the bath temperature of 32°C for 30 min with an ACSF flow rate of 1.2-1.4 ml/min. Extracellular DA concentration ([DA]_o_) was monitored with FCV at carbon-fiber microelectrodes (CFMs) using a Millar voltammeter (Julian Millar, Barts and the London School of Medicine and Dentistry). CFMs were fabricated in-house from epoxy-free carbon fiber ∼7 μm in diameter (Goodfellow Cambridge Ltd) encased in a glass capillary (Harvard Apparatus: GC200F-10) pulled to form a seal with the fiber and cut to a final tip length of 100-130 μm. The CFM was positioned 100-120 μm below the tissue surface at a 45° angle. A triangular waveform was applied to the carbon fiber scanning from −0.7 V to +1.3 V and back, against a Ag/AgCl reference electrode at a rate of 800 V/s. Evoked DA transients were sampled at 8 Hz, and data were acquired at 50 kHz using AxoScope 10.5 /10.7 (Molecular Devices). Evoked oxidation currents were converted to [DA]_o_ from post-experimental calibrations of the CFMs. Recorded FCV signals were identified as DA by comparing oxidation (+0.6 V) and reduction (−0.2 V) potential peaks from experimental voltammograms with currents recorded during calibration with 2 μM DA dissolved in ACSF.

DA release was evoked every 2.5 minutes with electrical or optical stimulation delivered out of phase with voltammetric scans. For electrical stimulation, a concentric bipolar stimulating electrode (FHC: CBAEC75) was positioned on the slice surface within 100 μm of the CFM. DA release was evoked using square wave pulses (0.6 mA amplitude, 2 ms duration) controlled by an Isoflex stimulus isolator (A.M.P.I., Jerusalem, Israel). For site sampling experiments with electrical stimulation, three stimulations were delivered at a given site before progressing to a corresponding site in the other slice. For optical stimulation of ChR2-expressing DA terminals, release was evoked using 5 ms pulses of 473 nm light delivered through a 10x water-immersion objective coupled to a CoolLED system (CoolLED pE-300; 26.6mW light power output) controlled by a Master-8 pulse stimulator (A.M.P.I., Jerusalem, Israel). Optically evoked DA events were evoked with 10 light pulses delivered at 25 Hz.

### Open Field Behavior

Behavioral testing was carried out during the dark phase of the light cycle under white lights. Male and female mice (P50-90) were habituated to the behavior room under a blackout curtain for 1 hour prior to testing. General locomotor activity and avoidance behavior was assessed using a 60 minute session in an open field chamber (40 cm length × 40 cm width × 34 cm height) made of transparent plexiglass (ANY-maze, Stoelting). Horizontal photobeams were used to measure rearing and were positioned at 9 cm above the chamber floor. The chamber was cleaned between each mouse with 70% ethanol followed by water and let dry for 5 minutes. The apparatus was cleaned with water and mild soap detergent at the end of each testing day. The experimenter was blind to genotype throughout the testing and scoring procedures.

For open field testing, the mouse was placed in the front, right hand corner of the chamber, facing away from the center and activity was recorded using both an overhead camera and side-facing camera. A smaller center field of 20 cm length x 20 cm width was defined digitally to measure the time spent in the center of the open field. The observer manually scored grooming bouts and time spent grooming during the first 20 min of the test. A grooming bout was considered any licking, nibbling, or scratching that lasted more than one second. If there was a break of more than 2 seconds between actions, this was considered a separate grooming bout. The following parameters were measured using ANY-maze tracking software: total distance traveled, time spent in center, and number of rears.

### Adipo-Clear brain clearing

Brains from male and female (P40-60) DAT-Flp;Nex-Cre;Ai65 mice were cleared using Adipo-Clear (29) and imaged as described previously (19). Mice were perfused with 20 ml of 1x PBS, followed by ice-cold 4% PFA (16% PFA diluted in 1x PBS, EMS – 15710-S), followed by 10 ml of 1x PBS. Brains were post-fixed in 4% PFA in 1x PBS with shaking overnight at 4°C. Brains were then washed for 3 × 1 h in 1x PBS with 0.02% NaN_3_ (Sigma Aldrich – 58032) at RT. The brains were then delipidated with the following steps. The samples were initially dehydrated with a MeOH gradient (20%, 40%, 60%, and 80%, 1 h each, Fisher – A412SK-4) with B1n buffer (in H_2_O – Glycine: 0.3 M, Triton X-100: 0.1% (v/v), NaN_3_: 0.01% (w/v) with pH adjusted to 7.0) shaking gently at RT. Samples were then washed 2 x 1 h in 100% MeOH followed by overnight incubation at RT in DiChloroMethane:MeOH (DCM, Sigma – 270997) 2:1. The next day, samples were washed for 1 h in 100% DCM followed by 3 washes in MeOH for 30, 45, and 60 minutes. Samples were then bleached in H_2_O_2_ buffer (5% H_2_O_2_ in MeOH; hydrogen peroxide 30%, Sigma – 216763) for 4 hours at RT. Samples were rehydrated in a reverse MeOH/B1n gradient (from 80%, down to 20%) at RT for 30 minutes each followed by a 1 h incubation with B1n buffer. Samples were then washed in 5% DMSO/0.3% glycine/PTxwH buffer (DMSO, Fisher – D128-4;Glycine, Sigma – G7126; PTxwH buffer: in PBS - Triton X-100: 0.1% (v/v), Tween 20: 0.05% (v/v), Heparin: 2 ug/ml, NaN_3_: 0.01% (w/v)). Samples were washed in PTxwH buffer 4x for 30 minutes, overnight, 1 h, and 2 h.

Brains were immunolabeled with anti-RFP primary antibody (final concentration 1:1000, rabbit, Rockland #600-401-379). The antibody was first diluted in PTxwH buffer (1:100 in 500 μl) then spun down (13,000 x g) and the supernatant was transferred to an additional 4.5 ml of PTxwH buffer in order to remove any debris. The final 1:1000 anti-RFP in PTxwH buffer was added to the brains and put on a shaker at 37°C for 11 days. Samples were then washed 2 × 1 h, 2 × 2 h, and 2 × 1 day in PTxwH buffer at 37°C. Secondary antibody was diluted in 500 μl of PTxwH and spun down at max speed for 30 minutes to remove any aggregates or debris. The supernatant (∼480 μl) was then added to 4.5 ml of PTxwH (final secondary concentration of 1:500) (secondary antibody: donkey anti-rabbit 647, Thermo Fisher Scientific – A-31573) and brains were incubated at 37°C with shaking for 8 days. The secondary antibody was then washed as described following the primary antibody.

Following immunolabeling, brains were dehydrated in an MeOH gradient with H_2_O (20%-80%, as before) shaking at RT for 30 minutes for each step. Samples were then washed 3x in 100% MeOH at RT for 30, 60, and 90 minutes followed by a wash with 2:1 DCM:MeOH overnight rocking at RT. The following morning, brains were washed in 100% DCM at RT 3x 1 h. The clearing step started with a wash in DiBenzylEther (DBE, Sigma – 108014) for 4 h, gently rocking at RT. Following this, the DBE was replaced with fresh DBE and brains were stored in the dark at RT prior to imaging.

### Light sheet imaging and whole brain projection analysis

Whole brains were imaged on a LaVision Ultramicroscope II lightsheet using a 2x objective and 3 µm Z-step size. Whole brain fluorescence was collected using 640 nm laser excitation and autofluorescence was collected using 480 nm laser excitation.

Image alignments between channels and sample auto-fluorescence to the Allen Brain Atlas (ABA) serial two-photon tomography reference brain, were performed using elastix (https://elastix.lumc.nl/). Computational identification, extraction, and skeletonization of axon collaterals and processes was performed using the U-Net-based 3D convolutional neural network TrailMap (19,42). Regional statistics were generated by transforming the arborizations into the Allen Mouse Common Coordinate Framework (CCF) using the same parameters and vectors from the registration step. Axon/process content in each region was normalized to the size of the region and the total projection content in each brain and was expressed as a density metric.

## Acknowledgements

This work was supported by an NIMH Brain Initiative grant and supplement to J.N. (U01 MH-105878, H.S.B. Co-Investigator) and a Brain Initiative R01 to L.L. (NS-104698). H.S.B. was supported by a fellowship from the Alfred P. Sloan foundation. H.S.B. and P.K. were supported by NARSAD Young Investigator Grants from the Brain & Behavior Research Foundation (#25073 to H.S.B. and #27458 to P.K.). H.S.B. and D.H. are Chan Zuckerberg Biohub Investigators. D.H. is a Pew-Stewart Scholar for Cancer Research supported by the Pew Charitable Trusts and the Alexander and Margaret Stewart Trust. D.H. is supported by the Siebel Stem Cell Institute. The NEX-Cre mice were kindly provided by Dr. Klaus-Armin Nave. We would like to thank Holly Aaron and Feather Ives in the UC Berkeley Molecular Imaging Center for their microscopy training and assistance. We thank Dr. Johannes De Jong and Dr. Stephan Lammel for their feedback on this project. We thank the rest of the Bateup lab for helpful discussions and comments.

## Supplemental Figures and Legends

**Supplemental Figure 1.**
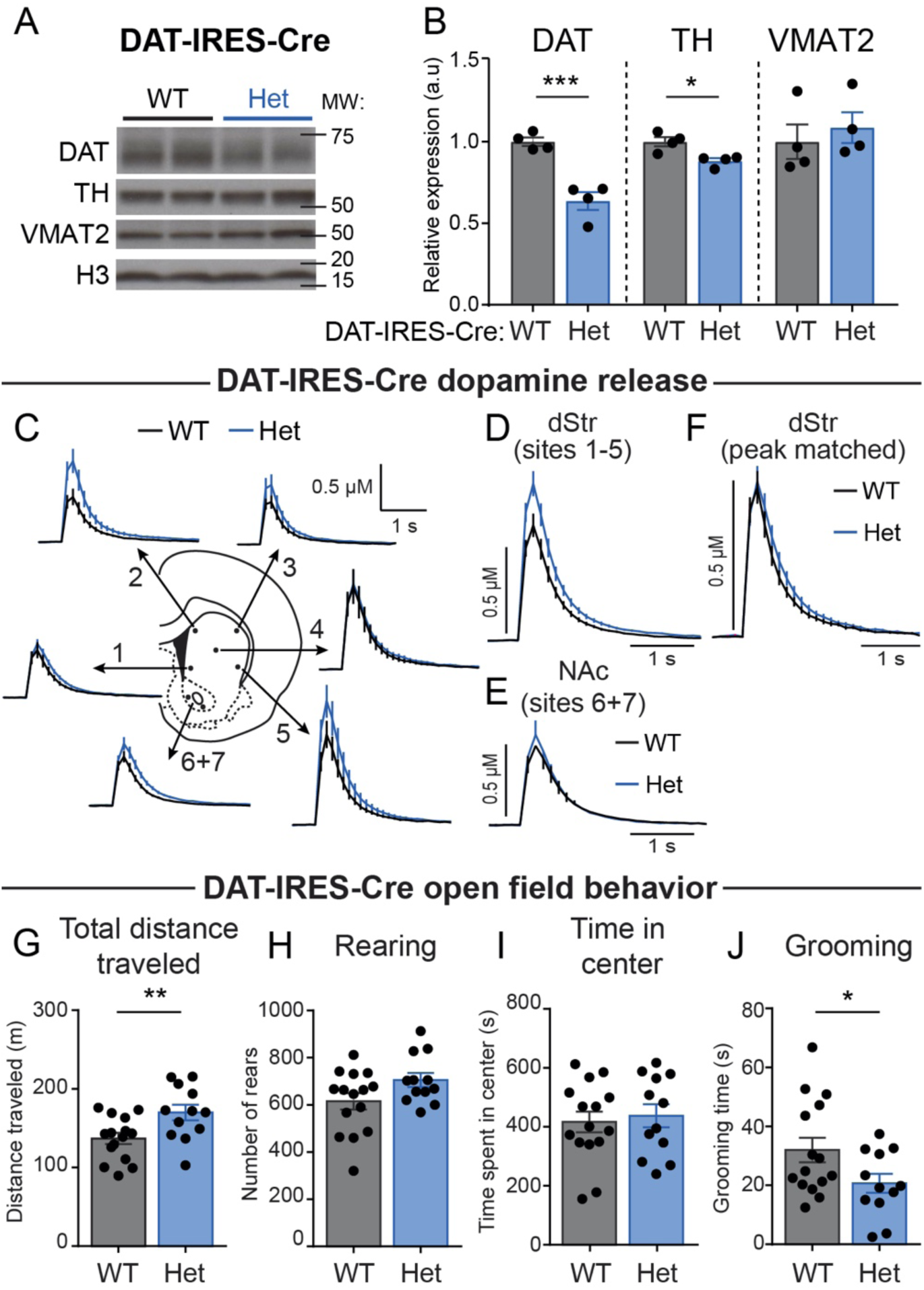
DAT expression and function in DAT-IRES-Cre mice (related to Figure 2). A) Representative western blot images of dopamine active transporter (DAT), tyrosine hydroxylase (TH), vesicular monoamine transporter 2 (VMAT2), and histone H3 (H3) loading control from striatal lysates from DAT-IRES-Cre wild-type (WT) and heterozygous (Het) mice. Two samples per genotype are shown. Blots were cropped to show the relevant bands. Molecular weight (MW) in KD is indicated on the right. B) Quantification of protein levels relative to histone H3, normalized to WT. Bars represent mean +/- SEM. Each dot represents the average of two striatal samples from one mouse (n=4 mice per genotype, 1 male and 3 females P100-120). DAT: p=0.0009, TH: p=0.0121, VMAT2: p=0.5655; unpaired t-tests. C) Fast-scan cyclic voltammetry (FCV) traces of extracellular DA ([DA]_o_) evoked by single electrical pulses in different striatal sub-regions in DAT-IRES-Cre mice. Traces are mean +/- SEM [DA]_o_ versus time (average of 27-28 transients per site from 4 pairs of DAT-IRES-Cre WT and Het mice, 1 male pair and 3 female pairs; P100-120). 1: ventromedial striatum, 2: dorsomedial striatum, 3: dorsolateral striatum, 4: central striatum, 5: ventrolateral striatum, 6-7: nucleus accumbens. D) Mean +/- SEM [DA]_o_ versus time (average of 80 transients per genotype) from the dorsal striatum sites #1-5. E) Mean +/- SEM [DA]_o_ versus time (average of 27-28 transients per genotype) from the nucleus accumbens sites #6-7. F) Region- and peak-matched mean +/- SEM [DA]_o_ versus time from the dorsal striatum of DAT-IRES-Cre WT and Het mice (average of 34-36 transients per genotype). p=0.072, single-phase exponential decay curve fit; WT tau=0.346, Het tau=0.355. G-J) Behavioral performance of DAT-IRES-Cre mice in a 60-minute open-field test. For all graphs, bars represent mean +/- SEM and dots represent values for individual mice. n= 15 WT mice; n=12 Heterozygous mice, all females; age P50-90. G) Total distance traveled in 60 minutes; p=0.0099, unpaired t-test. H) Total number of rears in 60 minutes; p=0.0668, unpaired t-test, I) Total time spent in the center of open field in 60 minutes; p=0.6976, unpaired t-test. J) Total number of grooming bouts in the first 20 minutes of open field test; p=0.0258, unpaired t-test.

**Supplemental Figure 2.**
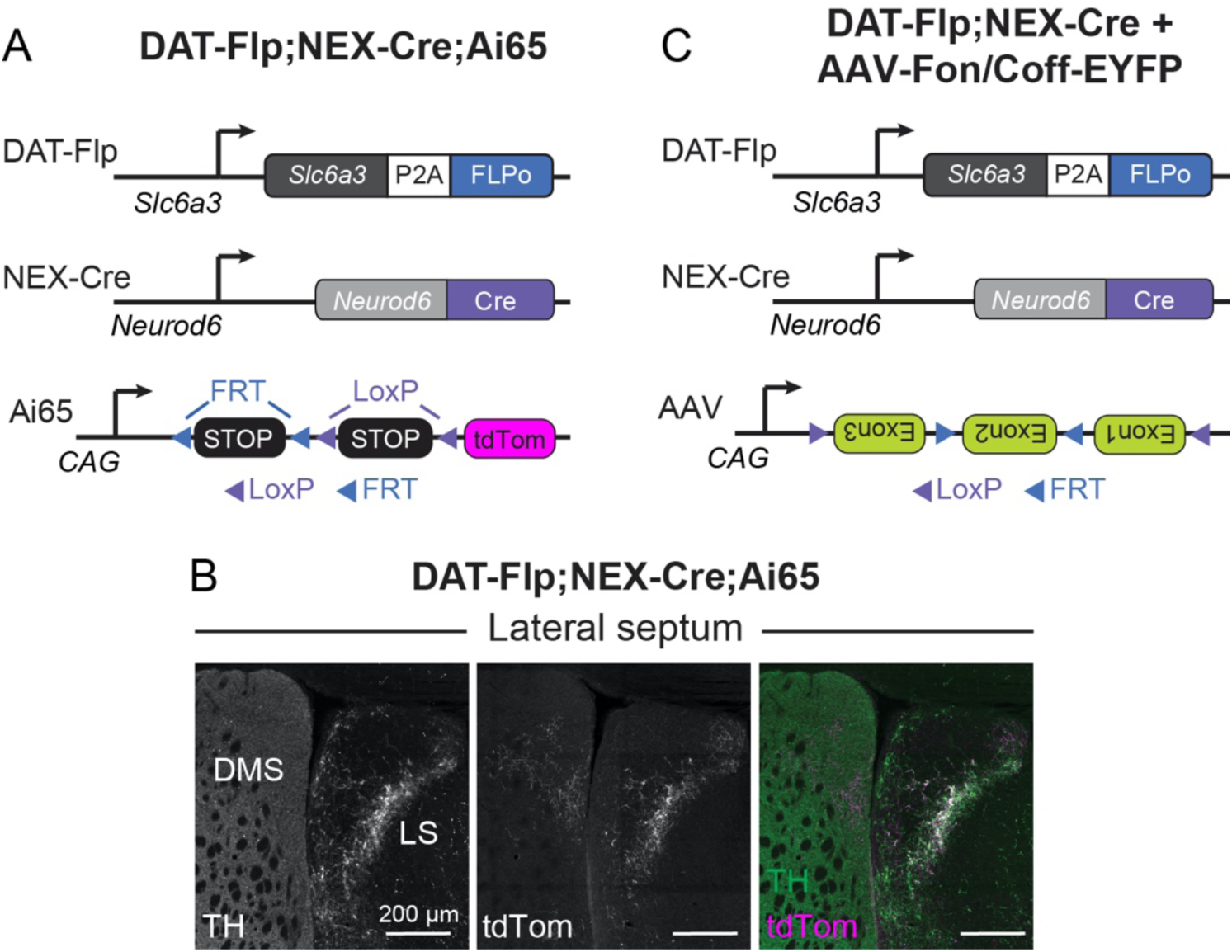
Intersectional labeling of neurons and projections in DAT-Flp;NEX-Cre mice (related to Figure 3). A) Schematic of the triple transgenic cross to generate DAT-Flp;NEX-Cre;Ai65 mice. In Ai65 mice, tdTomato is expressed in cells that express both Flp- and Cre-recombinase (30). B) Confocal images of projections from DAT-Flp;NEX-Cre;Ai65-expressing neurons to the lateral septum (LS). Left panel shows TH immunostaining. Middle panel shows tdTomato+ projections in the LS. Right panel shows a merged image of TdTomato (magenta) and TH (green) overlap in the LS. DMS = dorsomedial striatum C) Schematic of the strategy to label DA neurons that do not express *Neurod6* (see experiment in Fig. 3L,M). A Flp-on, Cre-off EYFP AAV construct was generated by Fenno et al (20) with artificially engineered introns to allow expression of full length EYFP only when Flp is expressed in the absence of Cre.

**Supplemental Figure 3.**
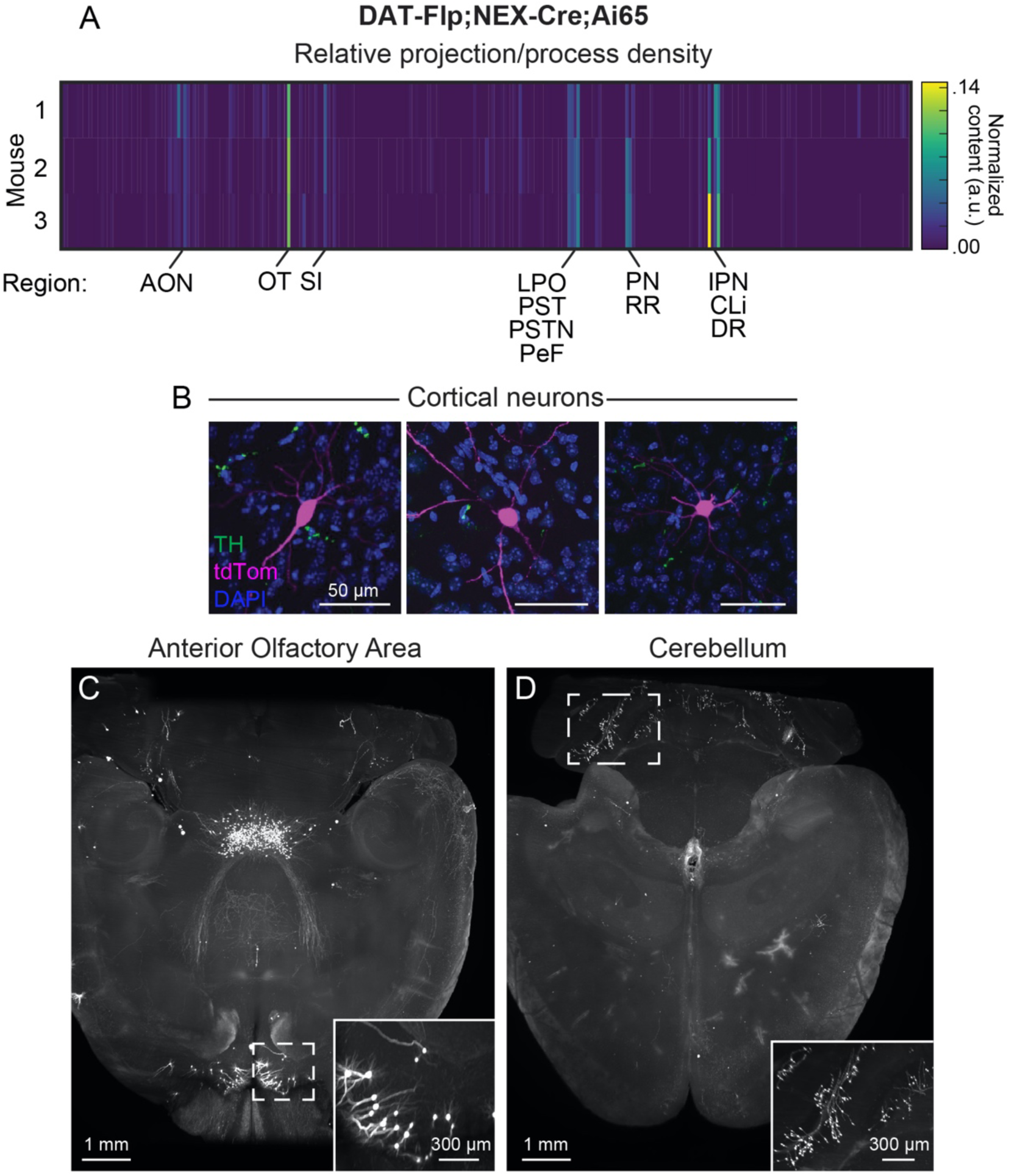
Whole brain imaging and projection analysis of DAT-Flp;NEX-Cre;Ai65 mice (related to Figure 4). A) Brains from 2 male and 1 female P120 DAT-Flp;NEX-Cre;Ai65 mice were processed with Adipo-Clear and imaged on a light sheet microscope. Projections and processes were quantified using TrailMap. Heatmap shows the total axonal/dendrite content of 277 brain regions using boundaries from the Allen Mouse Common Coordinate Framework (CCF). Values are normalized to both the region volume and total projection content per brain (taken from Supplemental Table 1 with ‘root’, ‘fiber tracts’, ‘ventricular systems’, and ‘Nucleus accumbens’ removed). The rows represent 3 independent brains. AON = Anterior optic nucleus; OT = Olfactory tubercle; SI = Substantia innominata; LPO = Lateral preoptic area; PST = Preparasubthalamic nucleus; PSTN = Parasubthalamic nucleus; PeF = Perifornical nucleus; PN = Paranigral nucleus; RR = Retrorubral area; IPN = Interpeduncular nucleus; CLi = Central linear nucleus raphe; DR = Dorsal nucleus Raphe. B) DAT-Flp;Nex-Cre;Ai65 mice exhibit tdTomato expression (magenta) in a small number of cortical neurons that do not co-express TH by immunostaining (green). C-D) Horizontal 450 μm Z-stack projections of light-sheet microscope whole brain images. DAT-Flp;NEX-Cre;Ai65 mice show consistent tdTomato expression in subpopulations of neurons in the anterior olfactory area (C) and cerebellum (D). See also Supplemental Table 1 and Supplemental Videos 1-3.

**Supplemental Table 1. Projection analysis of DAT-Flp;NEX-Cre;Ai65 mice using TrailMap.** Table of the TrailMap quantification of projection/process density in DAT-Flp;NEX-Cre;Ai65 mice. Listed are axon density values (arbitrary units) for the 3 measured brains and their average for the 281 regions from the Allen Brain Atlas. Also indicated are the Allen Brain Atlas ID numbers for each region and the abbreviation.

**Supplemental Video 1. 3D reconstruction of a single DAT-Flp;NEX-Cre;Ai65 brain.**

3D reconstruction of the raw signal from a single DAT-Flp;NEX-Cre;Ai65 mouse brain optically cleared using Adipo-Clear and imaged on a light sheet microscope.

**Supplemental Video 2. XZ-projection through TrailMap-extracted processes of DAT-Flp;NEX-Cre;Ai65 brains.**

Video of TrailMap-extracted processes from 3 DAT-Flp;NEX-Cre;Ai65 mouse brains overlaid on the Allen Mouse Common Coordinate Framework (CCF). Projections from the individual brains are color coded and the NAc region is marked off since the axons were too dense to consistently resolve in this region. Axons/processes are from 100 Z slices (5 μm/slice) and are contiguous rostral to caudal. The reference slice is a single slice from the middle position of the 100 Z slices.

**Supplemental Video 3. 3D rendering of TrailMap extracted processes in DAT-Flp;NEX-Cre;Ai65 mouse brains.**

3D rendering of TrailMap-extracted axons and dendrites from 3 DAT-Flp;NEX-Cre;Ai65 mouse brains overlaid on the Allen Mouse Common Coordinate Framework (CCF). Processes are color coded by mouse.

